# Reproducibility of biophysical *in silico* neuron states and spikes from event-based partial histories

**DOI:** 10.1101/2023.04.15.536945

**Authors:** Evan Cudone, Amelia M. Lower, Robert A McDougal

## Abstract

Biophysically detailed simulations attempting to reproduce neuronal activity often rely on solving large systems of differential equations; in some models, these systems have tens of thousands of states per cell. Numerically solving these equations is computationally intensive and requires making assumptions about the initial cell states. Additional realism from incorporating more biological detail is achieved at the cost of increasingly more states, more computational resources, and more modeling assumptions. We show that for both point and morphologically-detailed cell models, the presence and timing of future action potentials is probabilistically well-characterized by the relative timings of a small number of recent synaptic events alone. Knowledge of initial conditions or full synaptic input history is not a requirement. While model time constants, etc. impact the specifics, we demonstrate that for both individual spikes and sustained cellular activity, the uncertainty in spike response decreases to the point of approximate determinism. Further, we show cellular model states are reconstructable from ongoing synaptic events, despite unknown initial conditions. We propose that a strictly event-based modeling framework is capable of representing the full complexity of cellular dynamics of the differential-equations models with significantly less per-cell state variables, thus offering a pathway toward utilizing modern data-driven modeling to scale up to larger network models while preserving individual cellular biophysics.

## Introduction

Computational neuroscience research employs a variety of modeling strategies to simulate neurons and networks. One such strategy, conductance-based neuron modeling, uses systems of differential equations with terms that represent the electrical and biophysical properties of excitable neurons. Such models are fundamental building-blocks for bottom-up design approaches in the modeling of the nervous system. Algorithmic advances [1–3], parallelized computing [4–6], and modern computing hardware [7–10] have advanced our ability to expediently run conductance-based models, but scaling such simulations to efficiently engage in the phenomenological study of the brain remains a challenge. Network simulations of conductance-based neuron models are possible given sufficient compute resources; e.g. Traub et al. 2005 (3.5k neurons) [11], Migliore et al. 2014 (13k-120k neurons) [12], Potjans & Diesmann 2014 (80k neurons) [13], Markram et al. 2015 (31k neurons) [14], Yang et al. 2019 (1m neurons on specialized hardware) [15], Billeh et al. 2020 (230k neurons) [16]. However, simulating the collective 86 billion neurons of the human brain with morphological and biophysical detail requires orders of magnitudes more processing power than what is currently feasible.

One alternative to simulating large systems of differential equations is to use an event-based modeling framework. Event-based neuron modeling considers a neuron model that responds to input events (synaptic stimuli events) with output events (resulting action potentials). Modeling synaptic communication using only events can accelerate network simulation by decoupling the simulation of individual cells in a network, making it possible to simulate large neuronal networks in parallel [17]. Further efficiency at the cost of biophysical interpretability may be gained by replacing the calculus of the conductance-based models with a simplified rule, such as the analytical solution to the Integrate and Fire (INF) neuron model [18]. Simulation of event-based models advances with each event, rather than by a predetermined timestep. The resultant low computational burden means that INF neurons are often the first choice when simulating large networks to study network structure and network dynamics [19] but the simplified neuron model reduces the biological relevance in the context of living nervous systems.

The event-based framework need not, however, be limited to simplified neuron models. Here, we introduce the use of an event-based framework as a way to represent biophysically detailed models based on their input/output (I/O) relationships instead of their state variables. We demonstrate that the spiking neuron behaviors are reproducible with a strictly event-based system of a small number of the most recent stimuli. We model the responses of Hodgkin-Huxley (HH; [20]) neuron models with and without morphology as event-based functions that take in the timings of recent cellular events, the onset of synaptic inputs and output spikes, as input. We show that reconstructed spike times and cellular state variables for both point and morphologically detailed models are often highly constrained by a relatively small sampling of the recent event space.

## Methods

### Conductance-based neuron modeling

All continuous-time conductance-based neuron simulation was accomplished using the NEURON (version 8.2) simulation environment [21]. We simulated single compartment neurons with HH dynamics [20], whose synaptic inputs were parameterized to exhibit five distinct behaviors (Table 1). Four of these parameterizations — Base, LW, LT, and LWLT – combine low and high stimuli weight parameterizations with slow and fast synaptic time constant parameterizations.The fifth model was parameterized specifically to exhibit bursting behavior similar to type II bursting (Fig 1). Stimuli events trigger exponentially decaying synaptic currents implemented with NEURON’s ExpSyn mechanism. Without loss of generality, we define the occurrence of an output spike as the moment during which the membrane potential crosses 0 mV from below within the single compartment for point cells and within the axon segment adjacent to the soma for the morphologically detailed CA1 pyramidal cell model.

**Figure 1:**
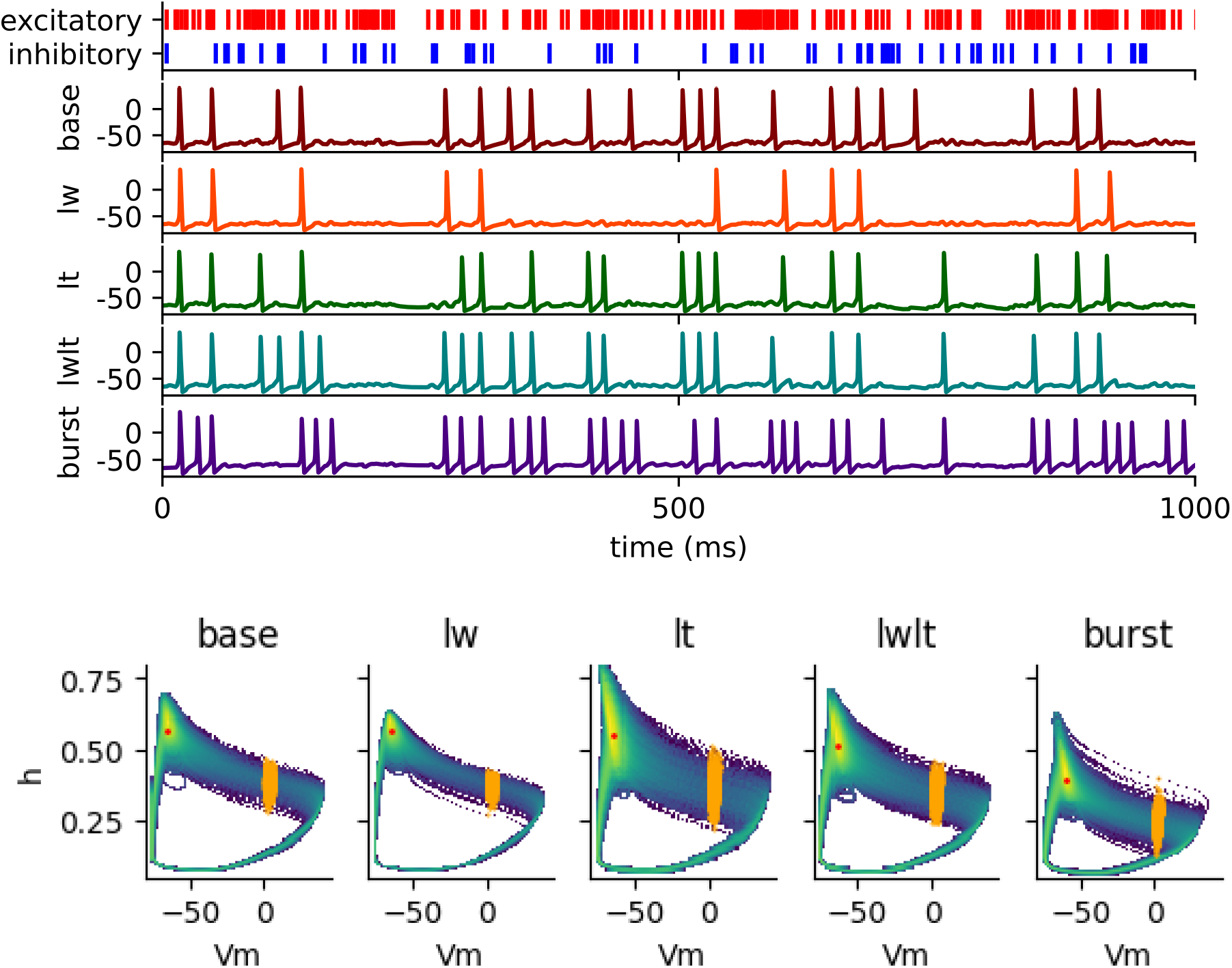
Point cell model responses. **Top**) Example membrane potential traces for the five synaptic parameterizations tested (Table 1) given the same excitatory and inhibitory stimuli demonstrate five different spiking behaviors, ranging from a low firing rate to the baseline behavior to a more bursty dynamic. **Bottom)** Density maps of the state space (*Vm* and *h*) of the five neuron types. Indicated are the models’ median state variable frame (red) and spiking state variable frames (orange).

**Table 1:** Synaptic connection parameterizations. We chose five synaptic connection parameterizations for single compartment cell models with HH dynamics to produce five distinct spiking behaviors that span short and long time dynamics, low and high reactivity to stimuli and spiking and bursting behavior.

|  | Excitatory synaptic weight | Inhibitory synaptic weight | Excitatory synaptic time constant | Inhibitory synaptic time constant | Average spike frequency | Coefficient of variation (CV) of ISIs |
| --- | --- | --- | --- | --- | --- | --- |
| Base | 0.0002 | 0.0005 | 2ms | 6ms | 22.14 Hz | 0.67 |
| Low Weight (LW) | 0.00015 | 0.0002 | 2ms | 6ms | 13.94 Hz | 0.80 |
| Long Tau (LT) | 0.0002 | 0.0005 | 10ms | 40ms | 17.62 Hz | 0.81 |
| Low Weight Long Tau (LWLT) | 0.00015 | 0.0002 | 10ms | 40ms | 23.10 Hz | 0.70 |
| Burst | 0.0001 | 0.0005 | 40ms | 20ms | 30.07 Hz | 0.84 |

For the studies incorporating morphology, we simulated a 3D CA1 pyramidal neuron [22,23] with HH mechanics in the soma and axon and leak channels in the dendrites. 10 excitatory and 5 inhibitory synapses were assigned at random locations on the cell’s dendrites in 30 different ways to account for spatial variability in stimulation. Each synapse received an independent stimuli stream generated with a Poisson process with an expected interval of 25 ms. We used random state variable frames from a 100,000 ms extended simulation as initializations to our experimental simulations to approximate the distribution of possible cellular states due to the unknown prior stimuli. For a given model and its synaptic parameterization, these include any of the 4 million state variable frames (*Vm*, *h*, *n*, and *m* for a point cell and *Vm*, *h*, *n*, and *m* for all compartments of the morphologically detailed cell) from the 100,000 ms extended simulation of that model. Spiking state variable frames are the subset of all frames which occur at the onset of an output spike (Fig 1 bottom). Median state variable frames represent the model’s median subthreshold state. To find the median state variable frames, we use the median value for each normalized variable and use the corresponding observed state variable frame with the minimum distance from those median values (red points in Fig 1 bottom). Here, each state variable is normalized separately.

### On-event simulation framework

To test the effect of varying event-based input encodings on neuron responses we developed an on-event simulation framework within NEURON that decouples event responses from simulation details like initial conditions, model states, and prior event history. This framework incorporates multiple parallel instances of the NEURON simulation environment of two varieties: one main simulation and multiple short-lived, on-event instances. The main simulation utilizes NEURON’s event driven framework [24] and only keeps track of the events (stimuli and output spikes); no latent state variables or membrane voltages exist in the main simulation. Stimuli events trigger parallel on-event functions which model the cell’s response given its recent event history and communicate the resulting events back to the main simulation. The response communicated is the next-spike-time (NST) — the simulation time until the next output spike event, which is set to null if the simulation is predicted not to spike provided no further input. Given a predicted real-valued NST, if the main simulation receives no other input before the predicted spike, it triggers and is recorded. If, however, a stimulus event occurs prior to the predicted spike, the spike is ignored, and the model’s behavior is recalculated using a new parallel on-event instance with the new recent event history.

### Limited event-based input encoding experiment

The limited event-based input encoding represents the total history of a cell as the timings of the last *n* events in the cell’s history. Incorporating this encoding in a neuron model constrains the total size of the input space (all historical events to *n* historical events) without changing the size of the response space (the NST).

We measured the error introduced by the limited event-based input encoding using a parallel instance of NEURON that runs a short conductance-based neuron model as our on-event function. These on-event instances are initialized with one of 10,000 random state variable frames. When the first event is an output spike, the set of random state variable frames used to initiate the on-event instance is constrained to a spiking state variable frame (Fig 1 bottom). On-event instances receive synaptic input corresponding to that of the recent event history and run for the duration of the recent event history plus 20 ms. The NST is the time delay from the last input event to the first spike in the remaining simulation; the first spike of the extended 20 ms; or null if there is no resulting spike. The on-event instance then communicates the corresponding NST to the main simulation which then proceeds with the next on-event instance.

We tested the degree to which the responses of both point cell and morphologically detailed cell models with HH mechanics change given limited event-based input encodings. We evaluate the effect of the limited event-based input encoding on neuron models in two primary contexts. First, we look at how the number of events (*n),* alters the presence and timing of single response spikes. To do so, we generated 1,000 different stimuli patterns with 30 combined excitatory and inhibitory stimuli. For each value of *n* in 3, 4, …30, we used the *n* most recent events of each of the 1,000 stimuli patterns in a simulation initialized with each of the 1,000 random state variable frames. The resulting 140 million simulations were used to analyze the robustness of the event-based framework to variations in the initial conditions given differing values of *n* inputs in the recent event history.

In this evaluation context we consider metrics to describe the consistency of spike presence, spike placement, as well as categorical descriptions of a response set describing its level of determinism. We introduce the metric of spike prediction coherence to measure the degree of consistency in the presence of spikes in the set of responses generated from randomized initializations. Given a stimuli pattern and *k* random state-variable frames as initial conditions, a fraction of their respective *k* simulations will respond with a spike. The spike prediction coherence measures the probability of pairwise agreement of spike presence given any two of the *k* simulations (ratio of spiking responses squared plus the ratio of non-spiking responses squared). To evaluate the evolution of responses observed as the recent event history changes, we define criteria to describe the spiking tendency of a set of responses as functionally deterministic or not. We consider a set of responses as deterministic non-spiking if less than 1% of the responses spike, as non-deterministic spiking if 99% of the responses spike and the standard deviation in spike timings is greater than 0.1 ms, as deterministic spiking if 99% of the responses spike and there is less than 0.1 ms standard deviation in the timings, and as non-deterministic if it meets none of the other criteria.

We also analyzed how this input encoding would affect the neuron model response over time. To do so, we first generated ground truth responses of the conductance-based models for comparison with simulations of the point and morphologically detailed neurons over a duration of 10,000 ms with Poisson generated excitatory and inhibitory input. Then, using the on-event simulation framework and the same input events, we ran extended simulations of the neuron models with the limited event-based input encodings. We ran these experiments using encodings with *n,* the number of input events in the encoding, in 3, 4, …, 30 with and without incorporating the timing of the last output spike as an input event in the encoding. Output spikes were modeled in the input encoding by initializing the simulation with a spiking history rather than with the median history, so the incorporation of the last output spike in the input encoding would truncate the encoding if there were stimuli events prior to the last output spike. With the resulting corresponding spike trains, we compared the observed continuous firing rates and interspike interval (ISI) distributions, as well as the distance between the two simulations’ spike trains calculated using the van Rossum metric [25] as implemented in the Elephant electrophysiology toolkit Python module [26].

### State variable reconstruction

We measured the ability to reconstruct the state variables of a simulation provided only the input events for each of the five synaptic parameterizations and the morphological CA1 pyramidal cell model. First, we recorded the state variables, input events and output events of a 100s simulation as a ground truth observation. 1,000 random spikes were chosen from the 100,000 ms simulations of each of the model types as starting positions for reconstruction. Since these reconstruction simulations begin at the instance of a spike, we initiated their state variables with a random spiking state variable frame of the respective model. The absolute error of the reconstructed state variables is the absolute value of the difference between the reconstructed state variable frame and the corresponding ground truth state variable frame.

## Results

To examine the viability of the on-event simulation strategy as an effective analog to conductance-based modeling, we ran three sets of analyses. First, we demonstrated that action potentials reset neuron states and act as natural barriers of information in neuron input history. Such information constraining events in the model act as evidence that the response of a cell is probabilistically determined by a recent subset of its total past input. We then showed that the specific historical information needed to constrain the possible responses is well characterized within the timings of the *n* most recent events. The specific value of *n* is model specific and generally tends to correlate with the magnitude of the time constants in the model. We found that models with a variety of different stimuli, like the varying placement of synapses along model morphology, benefit from categorically distinguishing these stimuli but do not show greatly increased demands in the total number of events needed despite their higher model complexity. Finally, we demonstrated the interoperability of conductance-based and event-based modeling by testing the extent to which the state variables of HH point and morphologically detailed models could be reconstructed from event timings only.

### Action potentials act of information bottlenecks in simulation

Action potentials in the HH system produce a stereotyped, cyclic pattern in its phase plane. The density around this phase plane is non-uniform, with depolarization displaying more variation and thus lower density than repolarization and refractory periods (Fig 1, Table 2). Because of their canonical shape, we considered action potential timings alongside stimuli timings as events in an event-based input encoding model.

**Table 2:**
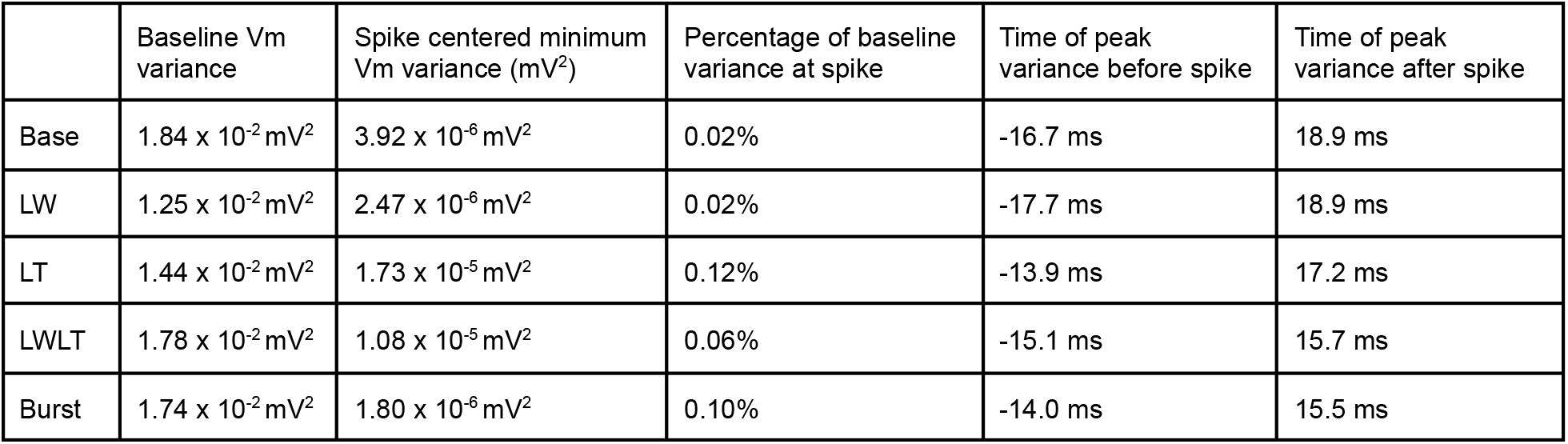
Comparison of membrane potential variance conditionally around spike events. Observed variance in membrane voltage of the entire simulation (baseline Vm variance) compared to that conditionally centered around the spike.

To test the extent to which action potentials constrain a cell’s state, we measured the spike-triggered-variance in the observed state variables. We found a consistent and large reduction in variance conditionally around spike events across all five of our HH model parameterizations when compared to the state variable variance observed across the entire simulation (baseline Vm variance) (Table 2). Models with longer time kinetics (LT, LWLT, and Burst) showed less of a reduction on the baseline variance (Table 2). The canonical form of the action potential resets the cells’ states and dramatically reduces the effects of previous stimuli on state variables (Fig 2).

**Figure 2:**
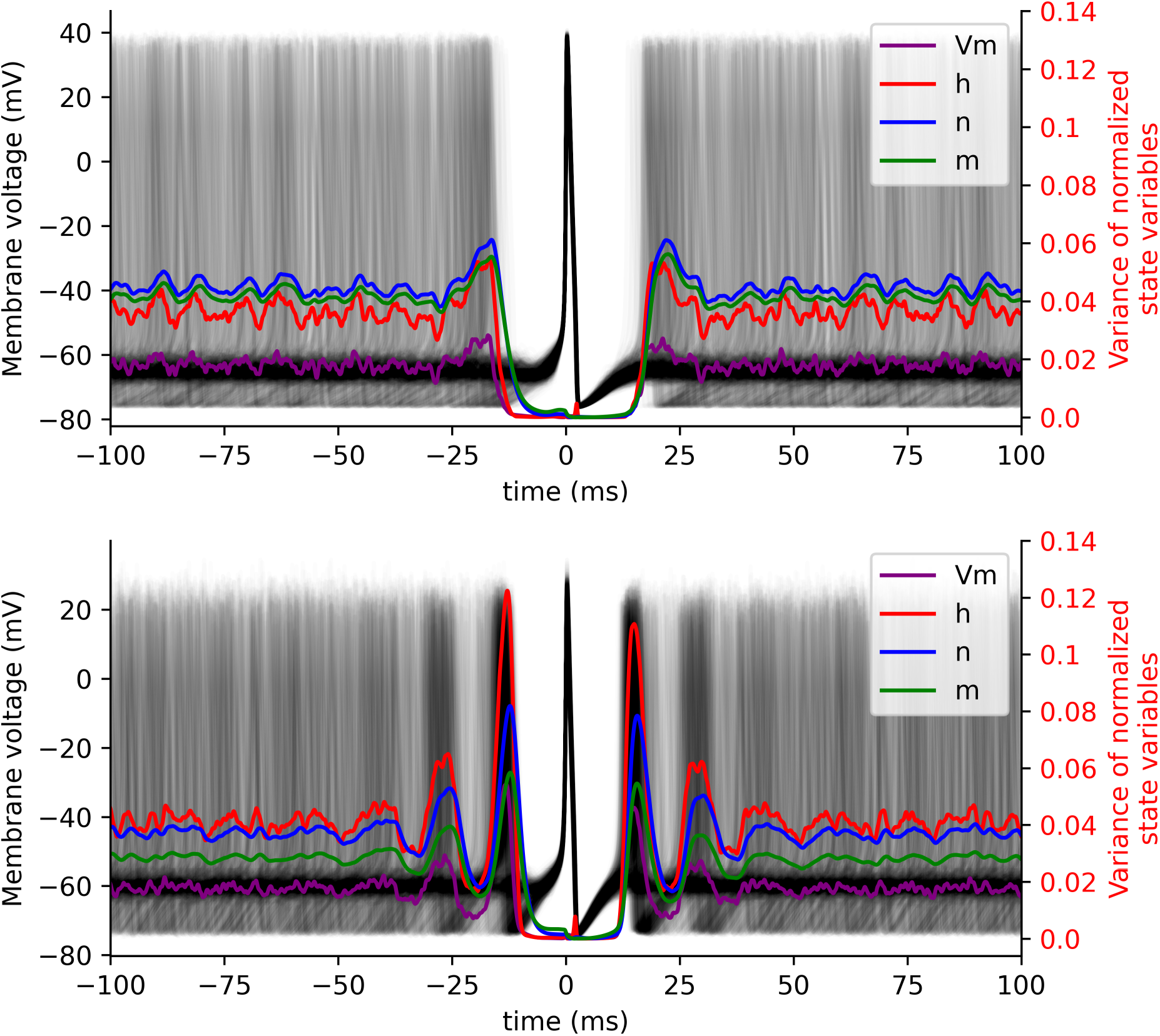
State space variation centered on spike events. 1,000 membrane voltage traces centered at a spike (black lines) and the observed variance in 40,000 simulation windows of normalized Vm (purple), h (red), n (blue) and m (green), for the spiking neuron (top) and bursting neuron (bottom). In both cases, the variance in state variables is significantly reduced relative to baseline for several milliseconds before and after the spike.

### Model determinism as a function of number of inputs in recent event history

As the number of inputs in the recent event history increases, the coherence of the responses increases and the NST standard deviation decreases: i.e. the distribution of responses tends towards determinism. This trend reaches a plateau after approximately 15 inputs for all synaptic parameterizations to varying degrees, with a considerable lag for the Burst parameterization (Fig 3). Likewise, the standard deviation in the timing of the observed spikes, NST standard deviation, approaches zero, but requires roughly 25 inputs in the recent event history, approximately 10 more input stimuli than the spike presence prediction (Fig 3). This indicates the placement of the spike timings is more sensitive to the exact state of the cell and requires more event history to accurately predict.

**Figure 3:**
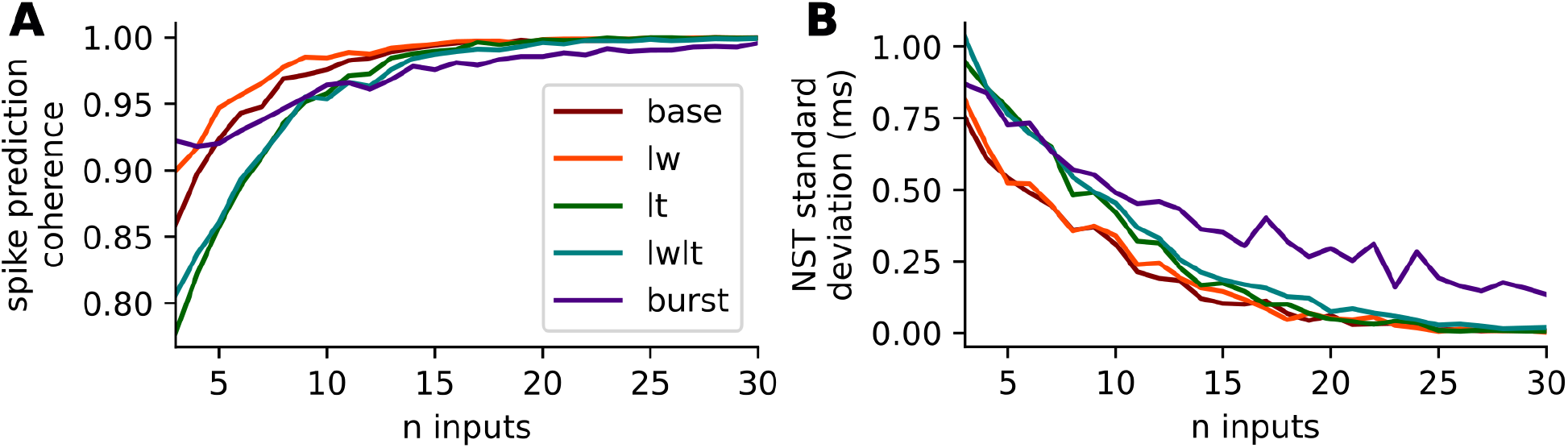
Spike prediction coherence and NST standard deviation as a function of size of recent event history. Two measures used to evaluate the determinism of spiking responses in a conductance-based Hodgkin-Huxley neuron model, given randomized initializations and recent event history. Spike prediction coherence **(A)**, is calculated as the ratio of spiking responses squared plus the ratio of non-spiking responses squared. NST standard deviation **(B)** is calculated from the distribution of observed timings of spiking responses across multiple trials. Both measures demonstrate generally increasing determinism with more inputs, with spike prediction coherence converging faster than spike timing.

The Base and LW synaptic parameterizations and the LT and LWLT synaptic parameterizations share roughly the same values for both the spike prediction coherence and the NST standard deviation, indicating that the time constant as the parameter with the most influence on the variance observed in the responses. The bursting parameterization requires the most event history to reach agreement on spike presence and more event history than our analysis included to reach agreement on spike placement. The plateau observed in both the spike prediction coherence and the NST standard deviation for all synaptic parameterizations indicate a high level of response determinism, and provide evidence that the model states are accurately encoded with the recent event history after 15-20 input events (Fig 3).

Investigating the evolution of responses as the number of events in the input encoding grows for a single stimuli pattern elucidates why the task of the precise number of events needed for response determinism is stimuli pattern specific. We show the responses for a specific input stimuli stream given a variety of initial conditions and an evolving number of stimuli in the input encoding as an example (Fig 4). Then, we characterized the distributions of responses given limited input encodings of a multitude of stimuli patterns as deterministic spiking, deterministic non-spiking, non-deterministic spiking, and non-deterministic and investigate the specific conditions in the stimuli patterns that force the response distributions to switch between these categories (Fig 5).

**Figure 4:**
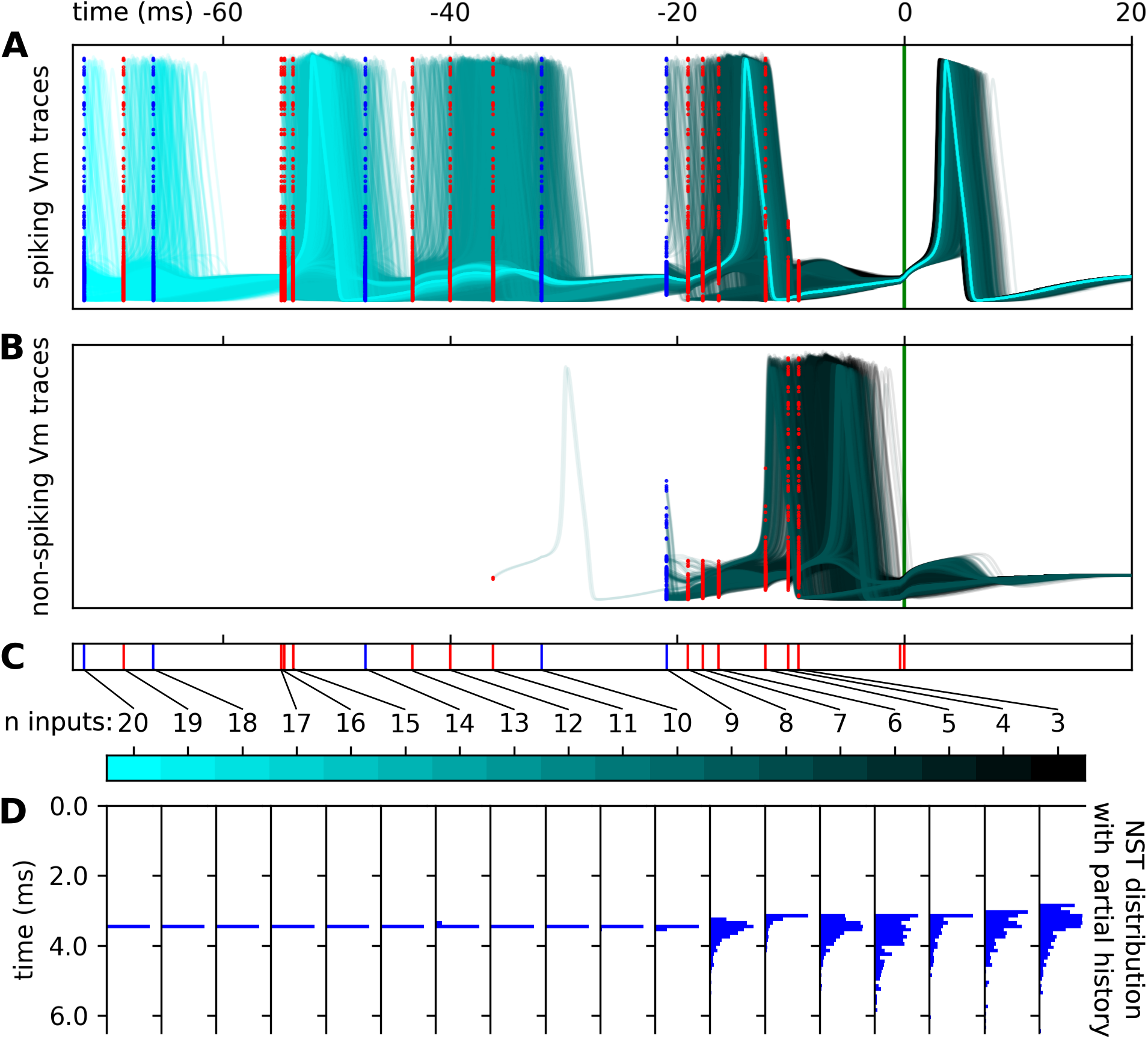
Evolution of spiking behavior as a function of number of inputs in input encoding for a given stimuli pattern. The Vm traces of spiking (A) and non-spiking (B) simulations with 20 (cyan) to 3 (black) input stimuli events in the recent event history for a single stimuli input pattern (C). Initializations of the simulations of these Vm traces indicated excitatory input stimuli events (red dots) and inhibitory input events (blue dots) correspond to the (C) input event raster. (D) The NST distribution corresponding to the specific stimuli events in (C) as histograms. The NST distribution partial history histograms are independently scaled for visual clarity, but have 0.67 spiking ratio for 3 inputs, 0.34 for 4 inputs, 0.29 for 5 inputs, 0.39 for 6 inputs, 0.78 for 7, 0.87 for 8, 0.82 for 9, and 1.0 for 10-20 inputs. This example highlights the specific stimulus instance responsible for collapsing the distribution of responses to a deterministic state.

**Figure 5:**
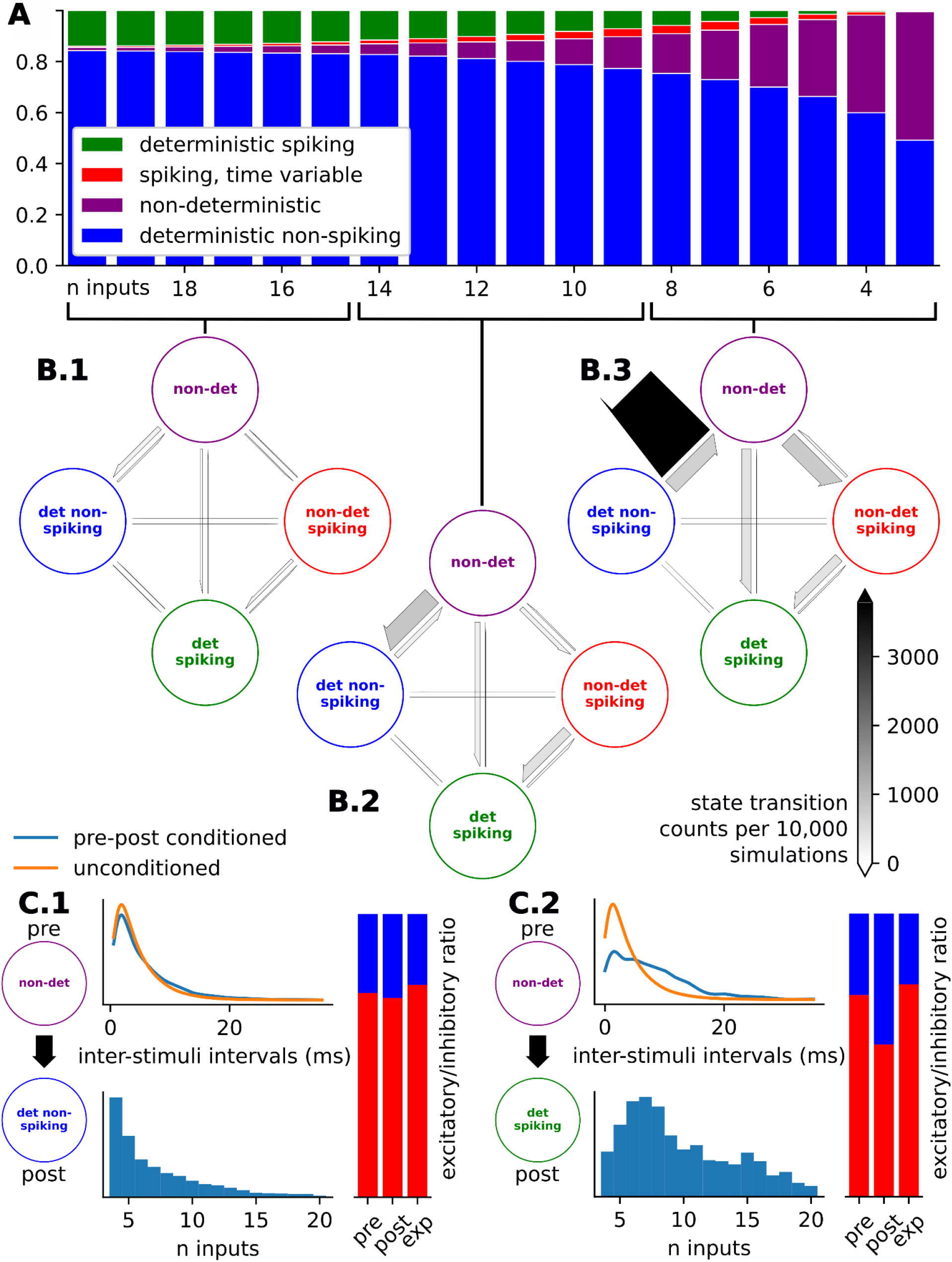
Response determinism of event-based input encodings with varying input history. **A**) Ratios of response distribution categories as the number of input events in the input encoding, *n*, increases shows a general increase in determinism when more inputs are known. **B)** Phase transition diagrams showing the transitions between response distribution categories for **B.1)** high values of n (n in 15, 16, … 20), **B.2)** medium values of n (n in 9, 10, … 14), and **B.3)** low values of n (n in 3, 4, … 8). **C)** Input conditions responsible for phase transitions between **C.1)** non-deterministic to deterministic non-spiking and **C.2)** non-deterministic to deterministic spiking phase transition. Input conditions include 1) inter-stimulus interval of the stimulus responsible for the transition (blue) compared to the unconditioned inter-stimulus intervals (orange), 2) histogram showing the relative presence of the specific transition category in relation to the number of input events, and 3) the ratios of stimuli types (excitatory: red and inhibitory: blue) of stimulus just before the transition (pre), of the stimulus responsible for the transition (post), and the expected, unconditioned stimuli type ratios (exp).

For the stimuli pattern in Fig 4, ambiguity on the presence of a spike persists up to the inclusion of the 10th input event, when the spike presence ratio goes from 0.82 (n = 9) to 1.0 (n = 10). This represents a switch from the category of non-deterministic to deterministic spiking, observed in the drastic change in NST distributions between the inclusion of this stimulus (Fig 4D). What is notable about the 10th input, the input responsible for this categorical change, is the large interval between it and the previous, 9th, stimulus. The tendency for long inter-stimuli-intervals corresponding to switches to the deterministic category is reflected in the statistical analyses showing the conditional difference in inter-stimuli-intervals for the stimuli responsible for non-deterministic to deterministic switching compared to the unconditional stimuli intervals (Fig 5C.2). Likewise, we notice that the response distributions switch fromon-deterministic to deterministic spiking conditionally more frequently when the post stimulus is inhibitory (Fig 5C.2).

The vast majority of deterministic spiking and deterministic non-spiking response distributions remain that way given as *n*, the number of input stimuli in the input encoding, increases. Further, the total number of transitions slows down as *n*, the number of stimuli in the input encoding, increases (Fig 5B). The observation that non-deterministic spiking response distributions tend to transition to deterministic spiking response distributions further emphasizes that spike timing prediction as a task requires more input information than spike presence prediction (Fig 5B).

### Event-based models with limited history reproduce realistic spike-trains

To evaluate the impact of limited input encoding in a full neuron simulation, we conducted a comparison of the observed spike trains and ISI distributions of the Hodgkin-Huxley (HH) model with and without the limited input encoding. When looking at the ISI distributions of the limited input encoding models that do not include the prior output spike as an input event, we noticed the emergence of unrealistic ISIs for low numbers of prior events (*n* < 5) suggesting this number of inputs is often incapable of accounting for the neuron’s refractory period (Fig 6). The occurrence of unrealistically small ISIs was greatly reduced when the input encoding included the timing of the last output spike as output spike events would inform the model of a refractory period (Fig 6). For large *n*, we found that including the output spike as a start point acts as a detriment to the replicability of the ISI distribution, especially for the synaptic parameterizations with long time constants as its inclusion often truncated the size of the input encoding (Fig 6).

**Figure 6:**
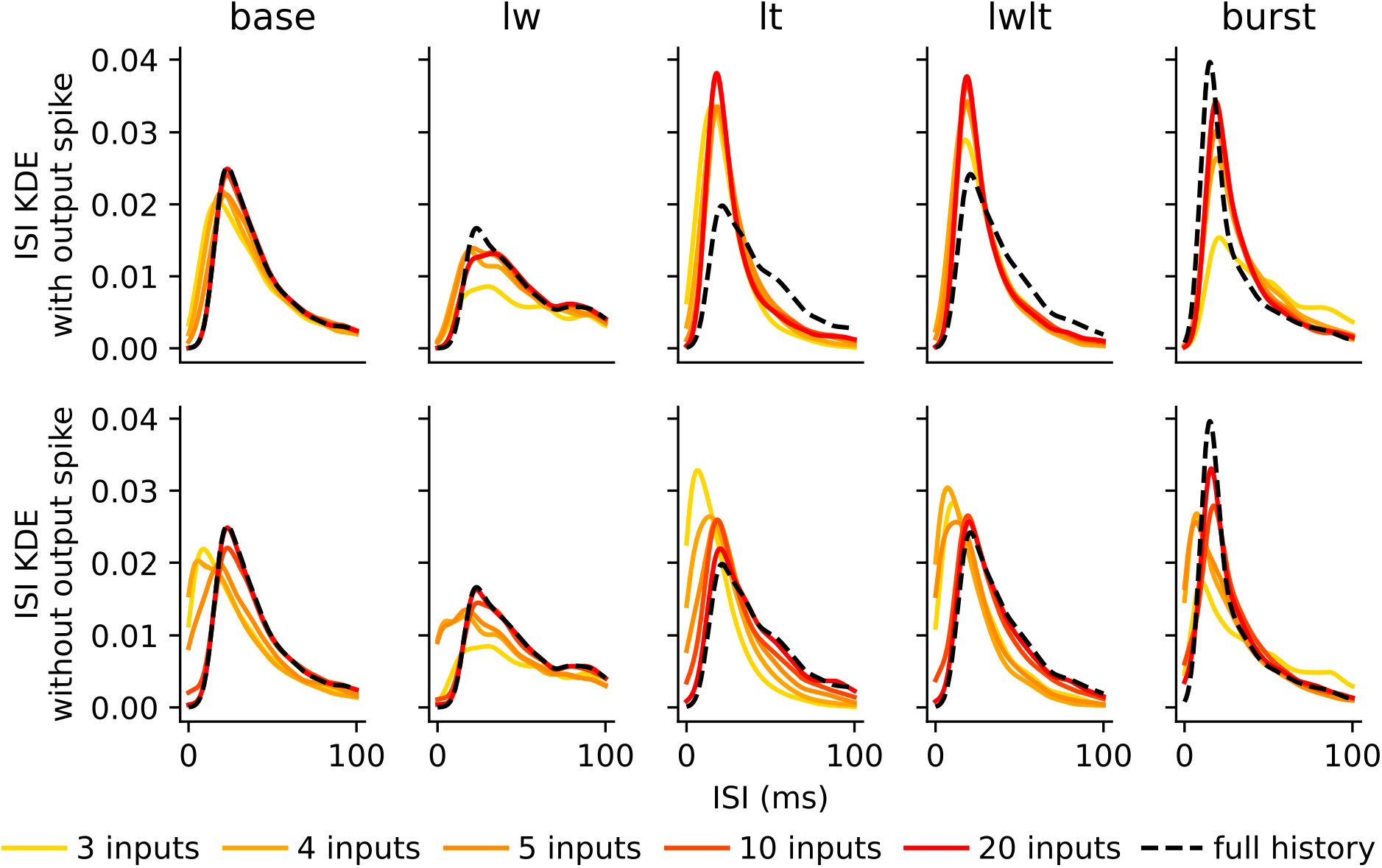
Evolution of ISI distributions as a function of number of inputs in input encoding: Observed inter-spike-interval distributions of spike trains with limited input encoding, shown as *n* increases, for the encodings with (**top**) and without (**bottom**) the inclusion of output events. These ISI distributions are compared to the ISI distribution of the HH model without the input encoding (full history). ISI distributions generally match the full history more closely with more input history. Including the time of the last output spike improved baseline distribution matching when few input times are known, but impaired the matching for the LT and LWLT cases when *n* is larger.

We compared the resultant simulated behavior of the neuron models with and without the input encoding using the van Rossum spike train edit distance metric [25] (Fig 7). The van Rossum distance decreased between the spike trains as the number of inputs in the encoding increased for all series, but converged to zero edit distance more quickly for the Base and LW parameterizations. The output spike train of the Burst parameterization proved to be the hardest to replicate, with a considerable van Rossum distance even after 40 input stimuli. For low numbers of n, the inclusion of the output spike in the encoding provides a more rapid reduction in van Rossum distance due to its elimination of unrealistically small ISIs (Fig 7), but plateaus after roughly ten inputs for the long time constant and bursting model parameterizations (Fig 7). In contrast, the output spike sufficiently filters prior input information for the Base and LW parameterizations, resulting in the performance of the encodings with and without the output spike converging to the same result. Including the output spike necessarily omits all prior synaptic information; we could not initialize the on-event simulations randomly and ensure a spike event at a given time. This constraint means that our analysis truncates stimuli history to the last output spike, meaning no further improvement is possible with additional stimuli events. The spike train distance results using the Victor-Purpura metric [27] were similar (not shown).

**Figure 7:**
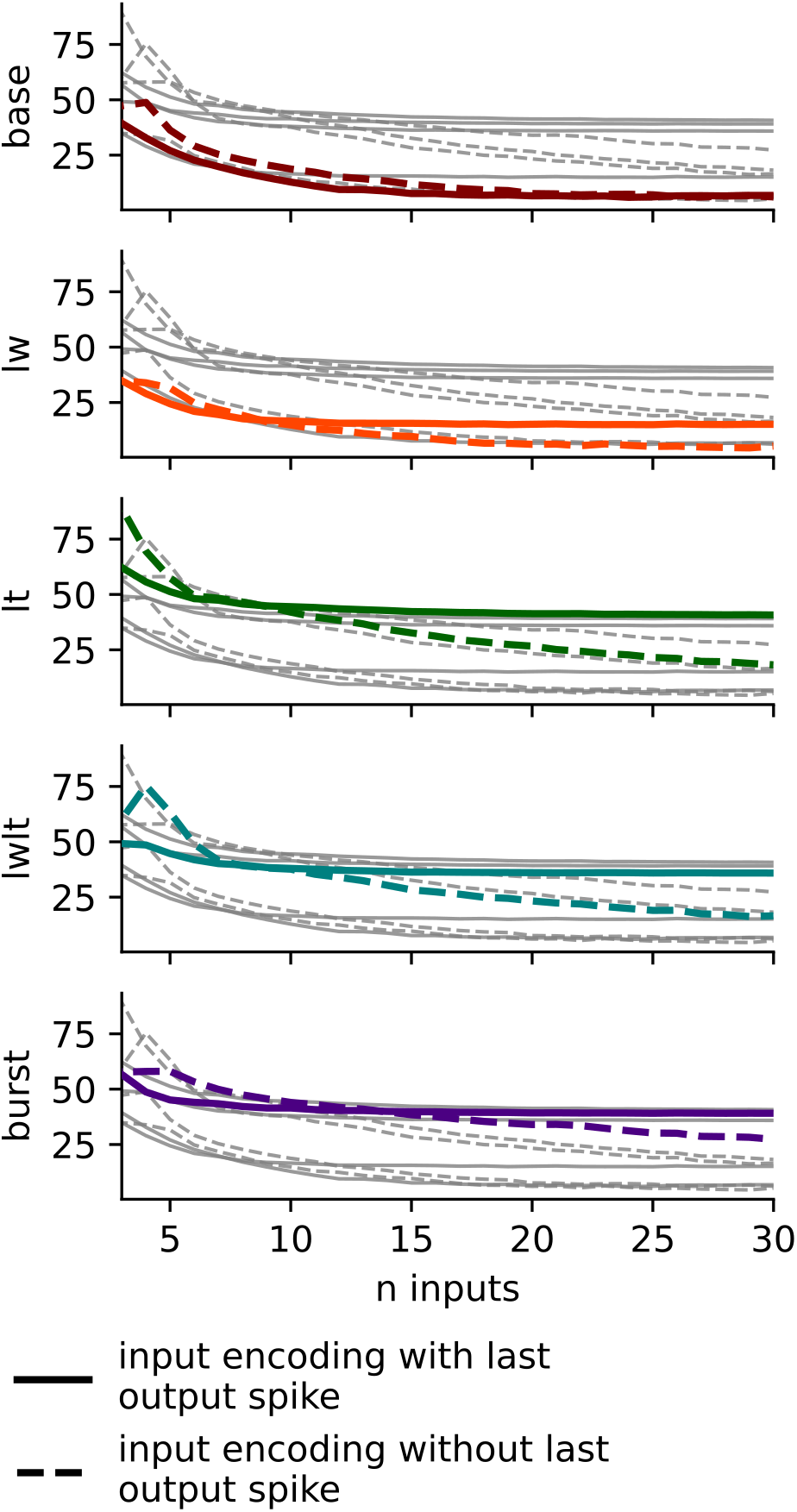
Spike train replicability of HH models with event-based input encodings as a function of number of inputs. van Rossum distance between observed spike trains from the neuron models with and without the event-based input encoding for each of the five HH model parameterizations. Shown are the distances for encodings which include the output spike (solid lines) and encodings which do not include output spike (hashed line). When very limited historical stimuli information is available, including the time of the last output spike reduced van Rossum error, but with larger amounts of stimuli history, for all but the baseline case the van Rossum error was lower on models generated without incorporating the last output spike time.

### Limited input encodings for morphologically detailed neurons requires categorically distinguish stimuli

Morphologically detailed neuron models such as the CA1 pyramidal cell used in this study [22,23] necessarily are more complex than their point cell counterparts. In addition to the multitude of compartments introduced by the model discretization each now with a volume and its corresponding set of state variables, morphologically detailed neurons can receive stimuli from a variety of different locations along the cell, each greatly influencing the behavior of the individual model. To account for this increased model complexity within the context of event-based modeling, the input encodings for morphologically detailed neuron models treat each distinct input location as an independent stimuli type in the same way the event-based point cell model distinguishes excitatory from inhibitory stimuli. Thus, the input-encoded morphologically detailed neuron with 10 unique excitatory and 5 unique inhibitory input locations receives the n most recent stimuli from all 15 sources.

In comparing the rasters of the ground truth response of the conductance-based morphologically detailed neuron to the event-based replications, we notice there are a few distinct ways the event-based system incorrectly predicts behaviors (Fig 8). First, when comparing the lowest values of n (*n* < 5) to the middle values of n (*n* in 5, 6, … 12), the event-based model switches from critically underestimating the presence of spikes to predicting too many spikes (demonstrated for emblematic example in Fig 8). This trend is evident in the van Rossum distances between the spike trains as the models with lower firing rates (cyan traces) have considerably lower distances for n in 3, 4, and 5 than those with higher firing rates (magenta traces) as they more closely match the firing rate (Fig 9). These disparate rates affect the general shape in the van Rossum distance to *n* inputs relationship as these low firing rate model’s (cyan traces) show an increase in the van Rossum distance between *n* of 3 and 5. The event-based models’ under-predicted spike presence for low values of *n* is more effective at predicting slower firing models. When the event-based model over-predicts spikes for *n* between 6 and 12, the output matches the spike trains of models with higher firing rates. Additionally, we notice the event-based system creates consistent errors over ranges of *n*. For instance, the event-based model incorrectly and consistently predicted the presence of spikes at 2,743 ms for values of *n* in 10-14 and at 3,451 ms for values of *n* in 5-17, among others (Fig 8). Despite this, the rasters mostly align and reach a consensus after the inclusion of roughly 15 input events (Fig 8), further evidenced in the plateau in the van Rossum distances plot (Fig 9). These results indicate that the responses of conductance-based morphologically detailed neuron models can be replicated with this limited input encoded event-based protocol.

**Figure 8:**
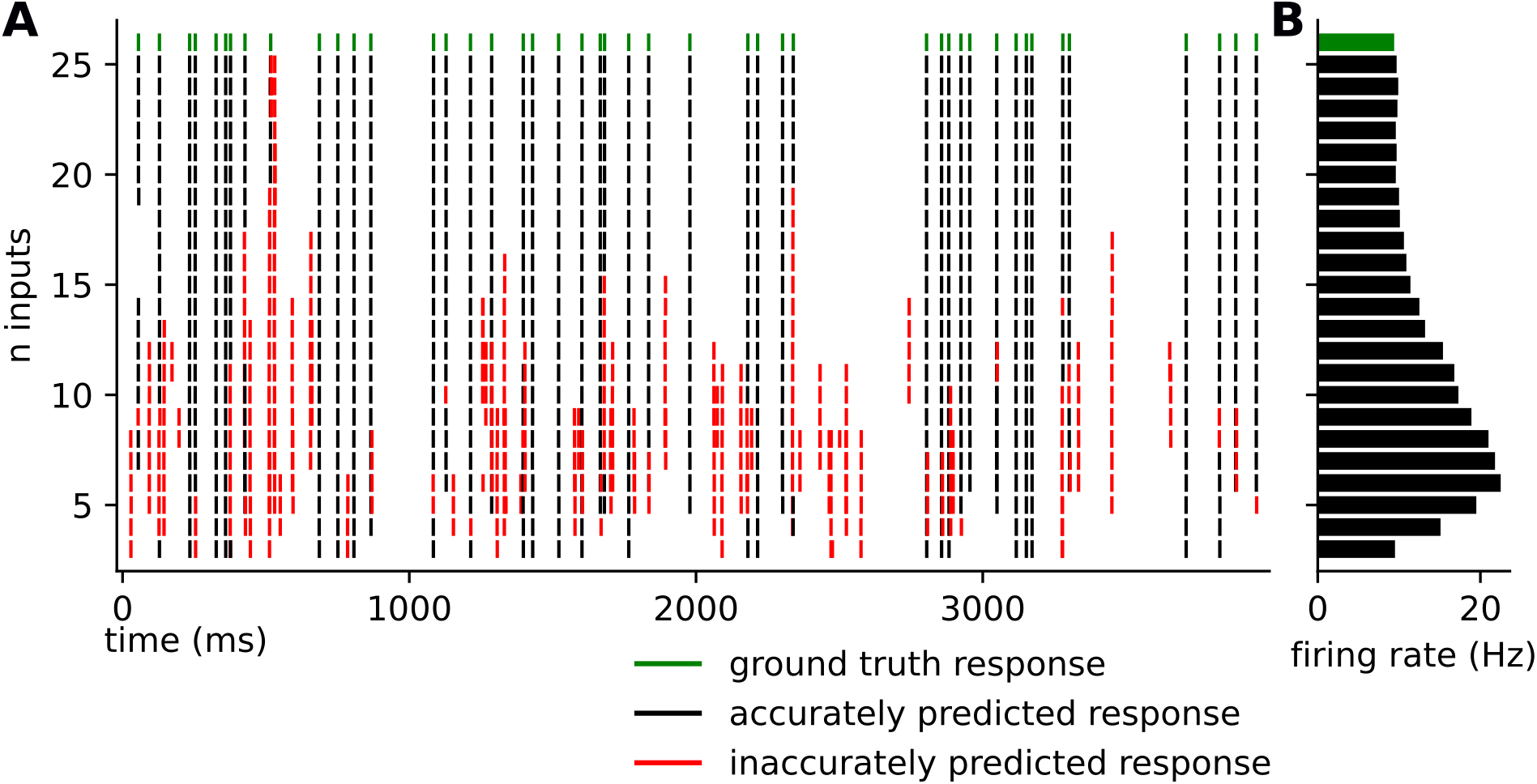
Evolving raster diagram of CA1 pyramidal cell outputs with varying recent event histories. **A**) Raster diagram showing an example of the simulated response of a morphologically detailed conductance-based CA1 pyramidal cell model as the ground truth (green) compared to the replicated responses of the event-based neuron model with recent event histories of size 3-25 (recent event histories of size 25-30 not shown because of minimal change). Black ticks indicate spikes within 1 ms of ground truth spikes, and red ticks indicate spikes > 1 ms away. Predicted spike times are broadly consistent when incorporating each additional input time, but become more accurate as more historical stimuli times are known. **B)** Corresponding firing rates of the ground truth ground truth spike train (green) and the replicated responses (black).

**Figure 9:**
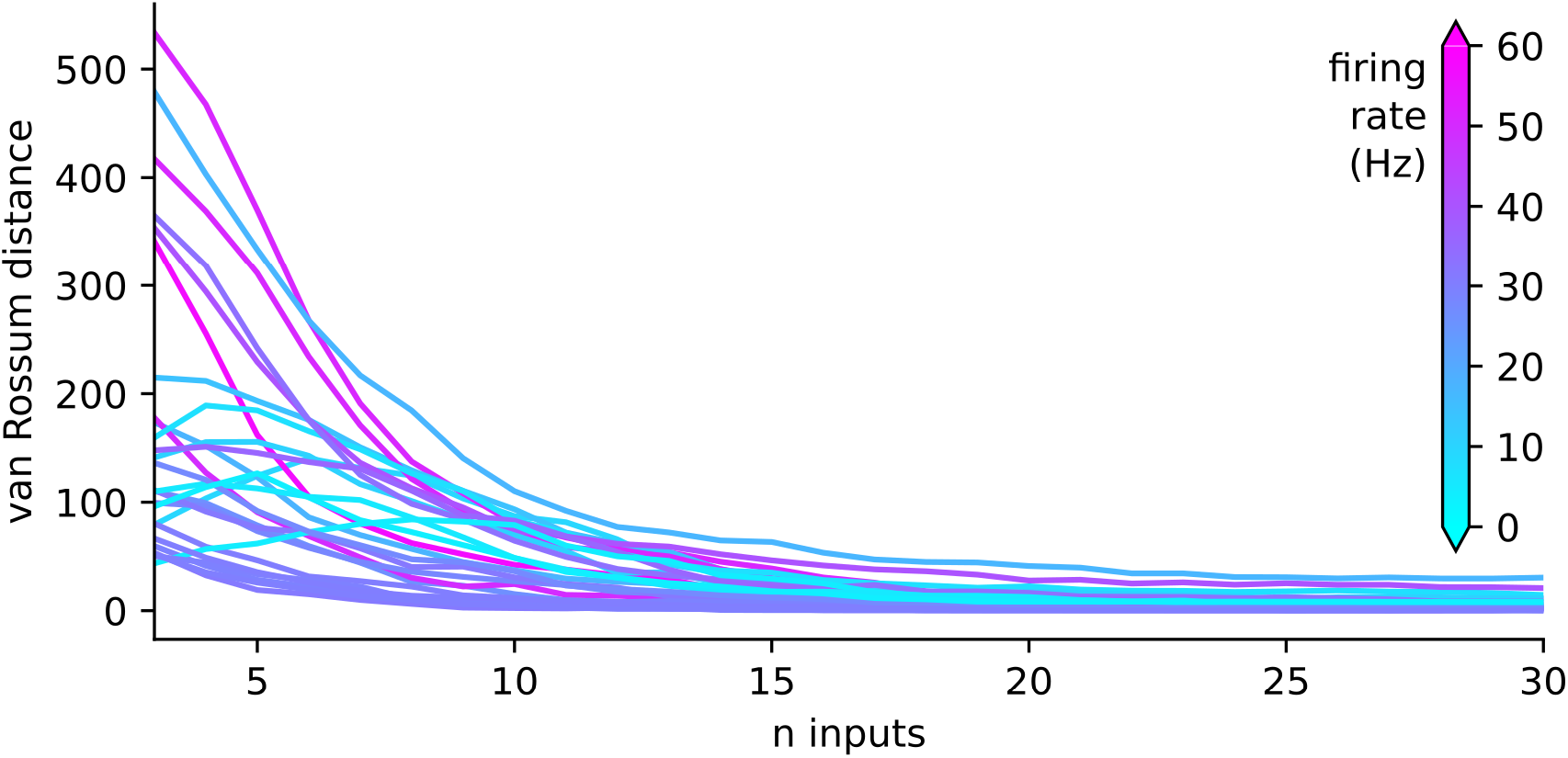
Spike train replicability of CA1 pyramidal cell model with event-based input encodings as a function of number of inputs. van Rossum distance between observed spike trains from the neuron models with and without the event-based input encoding for the morphologically detailed CA1 pyramidal neuron. Low firing rates show a slight increase in van Rossum error for the first few added inputs, but in general error decreased when additional input timings were included in the model.

### Reconstruction of state variables from a strictly event-based model

We reconstructed the continuous state variables of time windows from an extended HH point cell simulation using only the event-based stimuli timings to test our ability to retroactively reconstruct the state variables of a given window from the timings of input events alone. The mean absolute error (MAE) of these reconstructed windows decreases given a larger reconstruction window very quickly for the two synaptic parameterizations with small time constants and more slowly for those with larger time constants (Fig 10).

**Figure 10:**
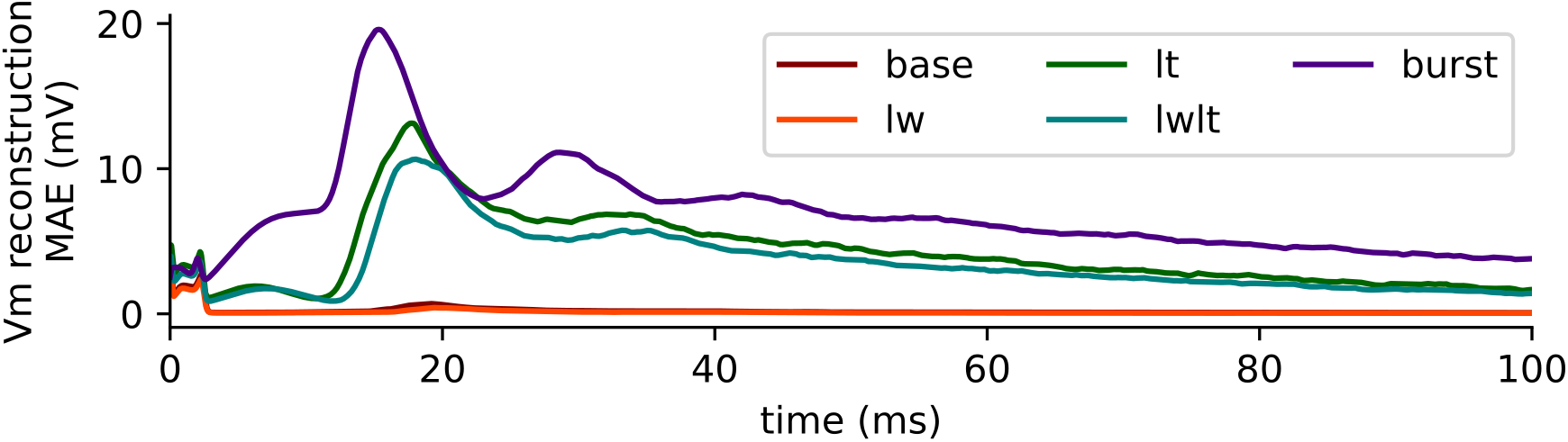
Reconstruction error as a function of time. Mean absolute error of membrane voltage reconstructions for each of the five synaptic parameterizations of the HH point cell.

We reconstructed the membrane potential for every segment in the morphologically detailed neuron for specific spike triggered windows of an extended simulation. When comparing the reconstruction MAE of different morphological regions of the model, we find the soma, apical and proximal dendrites behave similarly, showing a steep drop in reconstruction MAE after 2 ms. The reconstruction error of the axon, however, exhibits heightened error between 7 and 20 ms, which is not seen in the other morphological structures, likely due to the fact that the model contains HH mechanics only in the axon and soma (Fig 11A). Further, the relationship between the segments’ path distance and reconstruction MAE at both 10 and 25 ms appears to trend upward for the axon but downward for each of the individual dendrites (Fig 11B and Fig 11C). Most of the error is isolated to the axon and dendrites with inhibitory stimuli connections (Fig 12).

**Figure 11:**
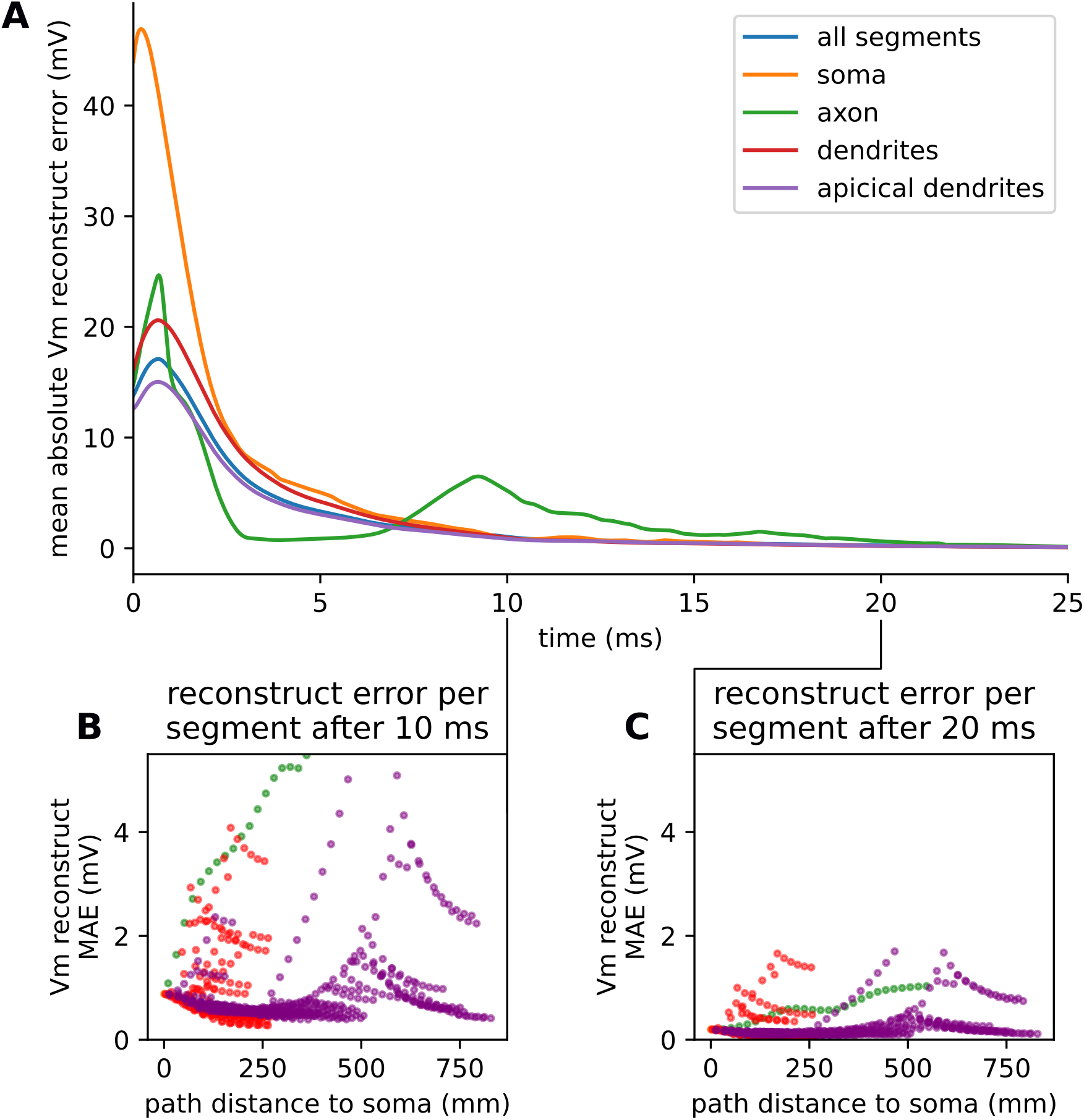
State variable reconstruction for an event-based morphologically detailed CA1 pyramidal cell model. **A**) Mean absolute error of membrane voltage reconstructions for each of the four neuron regions and their aggregate in the morphologically detailed CA1 pyramidal cell model. **B)** Scatter plot comparing the mean absolute membrane voltage reconstruction error and the path distance to the soma for each segment in the model at 10 ms. **C)** Scatter plot comparing the mean absolute membrane voltage reconstruction error and the path distance to the soma for each segment in the model at 20 ms.

**Figure 12:**
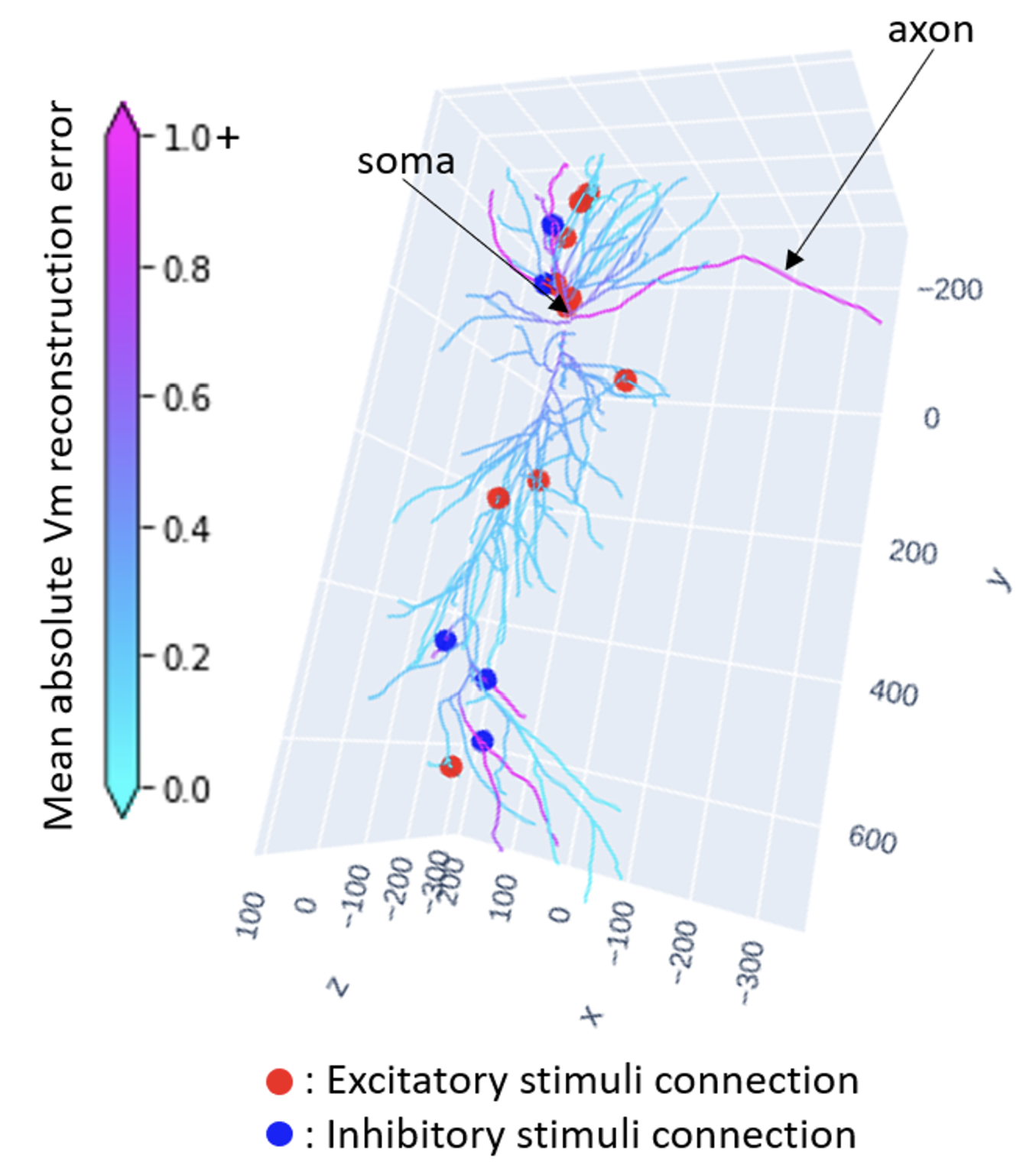
Reconstruction errors by segment after 20ms for an event-based morphologically detailed CA1 pyramidal cell model. 3D render of morphologically detailed CA1 pyramidal cell model with excitatory synapses marked with red dots, inhibitory synapses marked with blue dots, and the model segments colored with the mean absolute membrane potential reconstruction error, capped at 1.0 to highlight contrast (maximum MAE value is 1.94).

## Discussion

### Event-based approach to modeling electrophysiological dynamics

Conductance based neuron models represent the changing electrical states of cells using differential equations. Given a specific input stream, conductance-based models simulate, among other things, the membrane potential of the cell in such a way as to resemble the experimental recordings of biological neurons. In the wake of the ever-increasing data on ion channels and cellular morphology, the complexity of conductance-based biophysical neuron models has grown exponentially over the years. From the four state variables of the famous Hodgkin-Huxley model [20] to the cable theory models of dendrites using spatial compartments [28] to the routine use of morphologically detailed neuron models with upwards of 16,000 state variables each (e.g. [29]), computational neuroscience is consistently demanding more computational resources to meet its growing mechanistic complexity.

In this paper, we introduce an on-event simulation framework that decouples the model’s responses from preexisting model states and demonstrate the feasibility of interpreting conductance-based models within this framework. Here we model a neuron as a function whose total history passes through a narrow input encoding, a relatively small number of its most recent input events, to output a response. We test the extent to which the input encoding introduces error itself, separate from the specific modeling of the neuron response, by generating the response to the encoded input with a small instance of the conductance-based model in NEURON. But the framework is not limited to modeling responses in this way. Within this framework, the formalized mathematics and sub-threshold activity of the conductance-based model, or any model of neural response, can be folded into this on-event function without the need for solving state variables along a given timestep.

Our analysis shows that generating a neural response from a limited set of the neuron’s full event history does, to an extent, differ from the responses of the conductance-based version. Despite identifying conditions to minimize this error, we view it as a fundamental limitation of using this modeling approach to precisely replicate the explicit neural output of fully deterministic conductance-based neuron models. Even the precise outcomes of conductance-based models depend on the specific model of arithmetic used [30] and choice of numerical ODE solution procedure (e.g. Euler vs Crank-Nicholson vs Runge Kutta). Any model is necessarily a reduction of a physical system and we view the difference of responses of conductance-based and event-based modeling as one of the many possible sources of variation inherent to any choice of model in computational neuroscience. This being said, event-based modeling will benefit from a detailed understanding of how its neural responses differ from those of its conductance-based analogs and requires further investigation.

The response variation introduced by our on-event simulation approach can be measured and greatly limited. We show that, with sufficient history, the error between the conductance model and the event-based model decreases, indicating that the event-based input encoding conveys the necessary information to reproduce realistic, non-linear spiking responses. A full accounting of every prior event is not necessary to produce consistent responses. Instead, the spiking response of the HH system, with and without morphology, is probabilistically determined by a combination of a small number of recent synaptic events and the time of the cell’s last action potential, and its state variables are retroactively approximately reconstructable from a log of the neuron’s input events. In particular, the probability distributions for the presence and placement of a neuron model’s spikes grows tighter with more information about its stimuli history and its spiking history. This general convergence holds for point cells as well as for morphologically detailed cells, with only a slight increase in the necessary input events to account for morphology despite the multiple magnitudes difference in number of state variables. We found that slow dynamics required more history. For bursting neurons we achieved better results without explicitly considering the time of the last spike, presumably because the last spike alone gives no information about where in the burst we are and because of the truncation of prior history. When this distribution shrinks to the point of being functionally deterministic, the predicted response will no longer change given more information and the error cannot be reduced. We observed that this shift to determinism happens for the vast majority of input streams around the inclusion of 10 to 15 input events depending on the specific input pattern and its synaptic parameterization.

### Evaluation of the event-based neuron model requires consideration of multiple contexts

We identified three major contexts for which to investigate the implications of event-based modeling — how it could affect the presence and placement of singular spikes, how well it can replicate sustained activity of a simulation of a neuron, and what it could change at the network level.

At the single spike level, conceptualizing the neuron as a function that responds to a given input with a consistent output lends itself to evaluating alternatives as predictive models. Metrics like spike presence prediction accuracy and spike time prediction error would quantify how these two modeling schemas differ but would fail to statistically measure the ambiguity in the distribution of responses. Instead we used spike prediction coherence and standard deviation of NSTs to quantify the variance within a set of responses to the same input given differing levels of history. For an input stimuli pattern, if the discordance in responses is small enough, we are able to say the outcome is functionally deterministic and the history is sufficient to characterize the response to that particular input. Thus, we categorically describe the observed sets of responses as spiking non-deterministic (high spike presence coherence but high NST standard deviation), non-spiking deterministic, spiking deterministic and non-deterministic. With this categorization we investigated the specific conditions that cause the resulting responses to become deterministic. Our approach to evaluating the single spike context for event-based simulation achieves the task of measuring the viability of this simulation strategy without framing it as a strict prediction task.

However, the error measured for individual spikes does not necessarily translate to potential errors on sustained cellular activity because of the nonlinearity of the system and potential for compounding error. Thus we evaluated the model in the context of a neuron simulation over a given duration; that is, how well does the continued simulated response of the event-based neuron match that of the conductance-based neuron. We used the van Rossum spike train distance to represent a veritable ‘edit-distance’ between the conductance-based neuron and event-based neuron responses given identical input stimuli. Further, the interspike-interval distribution was used as a summary metric to denote large-scale differences in the overall behavior of the respective models.

Network level metrics like local field potential (LFP), synchrony, and firing rates would identify downstream consequences of the differences in responses of the conductance-based and event-based neuron models. However, we found that these metrics are too problem-specific and require additional modeling considerations, such as details on how the network is generated and connected. Controlling for these new modeling considerations would be challenging and limit the generalizability of this analysis to other neuron models. Additionally, neural coding research analyzes neuron and network models in mathematical and information processing contexts (e.g. [31–35]) that elucidate the detailed behavior of HH and other neural oscillatory systems under specific conditions. Our research, instead, aims to assess the feasibility of a generalizable modeling framework and assess how learnable the output responses of a single neuron is from the input. The data-driven nature of our evaluation makes it extendable to any spiking neuron model.

### Event-based framework as a bridge between mechanistic and phenomenological modeling

Event-based modeling is not new to network simulation; the use of INF neurons have paved the way for scaling up networks (e.g. [36]) and event communication is well integrated into NEURON [24] and other tools. What remains lacking is an efficient method to incorporate biophysical complexity at the individual neuron level for network simulations at scale. Given that input response of conductance-based and event-based neuron models are measurably congruent, we believe that event-based modeling, as shown in the present work, is a promising method for incorporating biophysical complexity at the individual neuron level in simulations at scale. Still, the specific types of error introduced by the event-based system may have unforeseen ramifications on the results of network simulations and remains a topic for further investigation [37].

Our event-based framework is not beholden to the specific input encodings we investigated here as it can incorporate any perceivable event of the model. For instance, incorporating the timing of the prior output spike in the encoding limited our ability to additionally consider stimuli events before it due to the specific way we set up our experiments in this study. This is an emergent limitation of the specific experiment we ran and is not the case for the framework as a whole. The full domain of applicability of this event-based approach remains to be discovered. Resources like ModelDB [38] provide large numbers of published cell models that could in principle be used for such a study; unfortunately, due to the heterogeneity of neuron models and the lack of a widely convertible model interchange format, the neuron models — even the ones using NEURON — are not readily adaptable to be incorporated in such a study.

The on-event modeling platform we built in NEURON for testing the predictability of spike response from limited stimuli history could, in principle, be used to introduce machine-learned event-based approximations to the biophysical models typically used in NEURON, either for individual cells or for entire networks. Machine learning has already accelerated network simulations [39], but relative to approaches that predict state variables with machine learning, an event-based model has the potential for further speed enhancements as it can skip from synaptic event to synaptic event rather than advancing through continuous time. Efficiently generating the training set and incorporating the effects of varying biophysical parameters of interest (e.g. synaptic conductances) are major challenges, but such rapid machine-learned models offer the potential of advance neuroscience simulation to incorporate more cells and longer run-time, bringing us closer to computationally connecting cellular dynamics to functional outcomes.

## Author Disclaimer

The content is solely the responsibility of the authors and does not necessarily represent the official views of the National Institutes of Health.

## Acknowledgments

We thank Dr. Ted Carnevale for sharing his insight and intuition in this discipline as well as J. NicFisk for their assistance in editing this manuscript. This work is supported by NIH 5T15 LM007056.

